# Benchmarking TCR-pMHC structure prediction: a unified evaluation and CDR3-based functional insights

**DOI:** 10.64898/2025.11.30.691400

**Authors:** Jiadong Lu, Xinyuan Zhu, Xinting Hu, Cheng Zhang, Fuli Feng

## Abstract

Interactions between T cell receptors (TCRs) and peptide-major histocompatibility complexes (pMHCs) are central to adaptive immunity. Recent advances in structure prediction tools have enabled atomic-level modeling of TCR-pMHC interactions. However, the lack of systematic evaluation forces practitioners to invest substantial resources in selecting appropriate tools. Here, we present a comprehensive benchmark of TCR-pMHC structure prediction with 70 previously unseen complexes and ten models spanning MSA-based, PLM-based, and docking-based approaches, revealing the superior modeling accuracy and docking quality of MSA-based methods, especially AlphaFold3. To further enhance the utility of AlphaFold3 predictions, we identify the pLDDT score of the TCR CDR3 region as an informative indicator of both structural correctness and functional relevance. Specifically, it enables up to 4.3% Top-1 success gain through reranking and captures mutation-induced affinity changes in over 80% of cases. Overall, our analysis would facilitate the practical usage of immune structure prediction models and guide the advancement of these models.

## 1 Introduction

The adaptive immune system, critically orchestrated by T cells, serves as a frontline defense against an array of threats, from rapidly evolving pathogens to cancerous cells [1, 2]. At the core of this defense, T cell receptors (TCRs) recognize antigenic peptides presented by major histocompatibility complex (MHC) molecules. These interactions form TCR-pMHC complexes that govern the specificity and breadth of immune responses. Understanding the structural basis of TCR-pMHC interactions is essential for uncovering the principles of immune recognition [3, 4] and for enabling applications such as vaccine development and personalized immunotherapy [5, 6]. However, experimental determination of TCR-pMHC complex structures remains challenging, due to the highly promiscuous nature of TCR-pMHC binding and the resource-intensive demands of structural biology techniques [7, 8].

In recent years, AlphaFold [9–11] has made significant breakthroughs in modeling protein-peptide [12], antibody-antigen [13–15], and other complexes [16], offering immense potential for its application in TCR-pMHC complex modeling. Building on AlphaFold’s success, a growing number of methods have emerged. These can be broadly categorized into: **multiple sequence alignment (MSA)-based folding methods** that leverage co-evolutionary information from large genetic databases; **protein language model (PLM)-based folding methods** that learn evolutionary features directly from single sequences, enabling faster predictions for fast-evolving proteins [17]; and **docking-based methods** which integrate deep learning-based folding models with physics-based docking algorithms. Recent approaches further incorporate acceleration strategies to meet the growing demand for high-throughput and rapid modeling [18]. However, existing studies often adopt inconsistent benchmarks and evaluation pipelines, which limit the fairness and practical applicability of current methods and make it difficult for practitioners to identify suitable tools.

Given the absence of consistent benchmarks for TCR-pMHC modeling, we systematically compare the three major categories of structure prediction methods (i.e., MSA-based, PLM-based, and docking-augmented) on 70 diverse and previously unseen TCR-pMHC complexes curated from TCR3d [19, 20]. The benchmark covers both class I (56 samples) and class II (14 samples) complexes. For each method, we evaluate docking quality using DockQ [21] and structural accuracy using RMSD and TM-score [22]. Our evaluation reveals that MSA-based methods, particularly AlphaFold3, consistently outperform all alternatives across both docking and structural metrics, achieving a median DockQ score above 0.6 for both class I and class II complexes. While AlphaFold3 delivers the best overall performance and strong potential for modeling immune recognition, AlphaFold2 remains competitive, and its efficient variants, TCRmodel2 [23] and ColabFold [18], achieve over 70-fold acceleration without compromising accuracy, enabling scalable high-throughput TCR-pMHC screening.

Building on the superior performance of AlphaFold3, we then explore its interpretability and practical utility by analyzing the predicted local distance difference test (pLDDT) score for the complementarity-determining region 3 (CDR3), which is a highly variable region that governs antigen-specific recognition and plays a decisive role in TCR-pMHC binding [24–27]. As a result, we find that AlphaFold3’s CDR3 pLDDT strongly correlates with both local modeling accuracy and overall docking quality. Leveraging this insight, we introduce a simple yet effective reranking strategy based on CDR3 pLDDT, which improves the Top-1 success rates of high- and medium-quality AlphaFold3 predictions by 2.9% and 4.3%, respectively. To further assess whether CDR3 pLDDT can aid AlphaFold3 in capturing the structure-function relationship, we model both wild-type and single-residue mutants and show that CDR3 pLDDT changes closely track mutation-induced binding affinity shifts, accurately reflecting functional alterations in over 80% of cases. These results demonstrate that CDR3 pLDDT not only reflects structural reliability, but also serves as a built-in signal that empowers AlphaFold3 to infer functionally meaningful changes in TCR-pMHC interactions.

In summary, we present a comprehensive and unified benchmark for TCR-pMHC complex modeling, covering diverse structure prediction paradigms and providing systematic evaluations of both accuracy and efficiency. Beyond the benchmark, we reveal the practical utility of AlphaFold3’s CDR3 pLDDT score for output selection and mutation effect prediction. Together, these contributions lay a scalable and interpretable foundation for TCR-pMHC modeling, with broad implications for immune repertoire analysis, therapeutic discovery, and personalized immunotherapy design.

## 2 Results

### 2.1 Benchmarking structure prediction methods for TCR-pMHC complexes

We began by benchmarking representative structure prediction models on a curated set of unseen TCR-pMHC complexes. Three categories of models (Fig. 1c) were evaluated: MSA-based folding models (AlphaFold2, AlphaFold3, Chai-1-MSA, RoseTTAFold2), PLM-based folding models (Chai-1-PLM, ESM3, ESMFold, OmegaFold), and docking-based approaches (AlphaRED and HDOCK applied to AlphaFold2 outputs). Our evaluations focused on the primary interacting regions of the complexes, including the variable domains of the TCR *α* and *β* chains, the peptide, and the peptide-binding domains of the MHC (Fig. 1a). The benchmark comprised 70 non-redundant structures (56 class I and 14 class II) from the TCR3d database (Fig. 1b), all released after September 30, 2021, ensuring they were excluded from the training data and structural templates of all evaluated models.

**Fig. 1.**
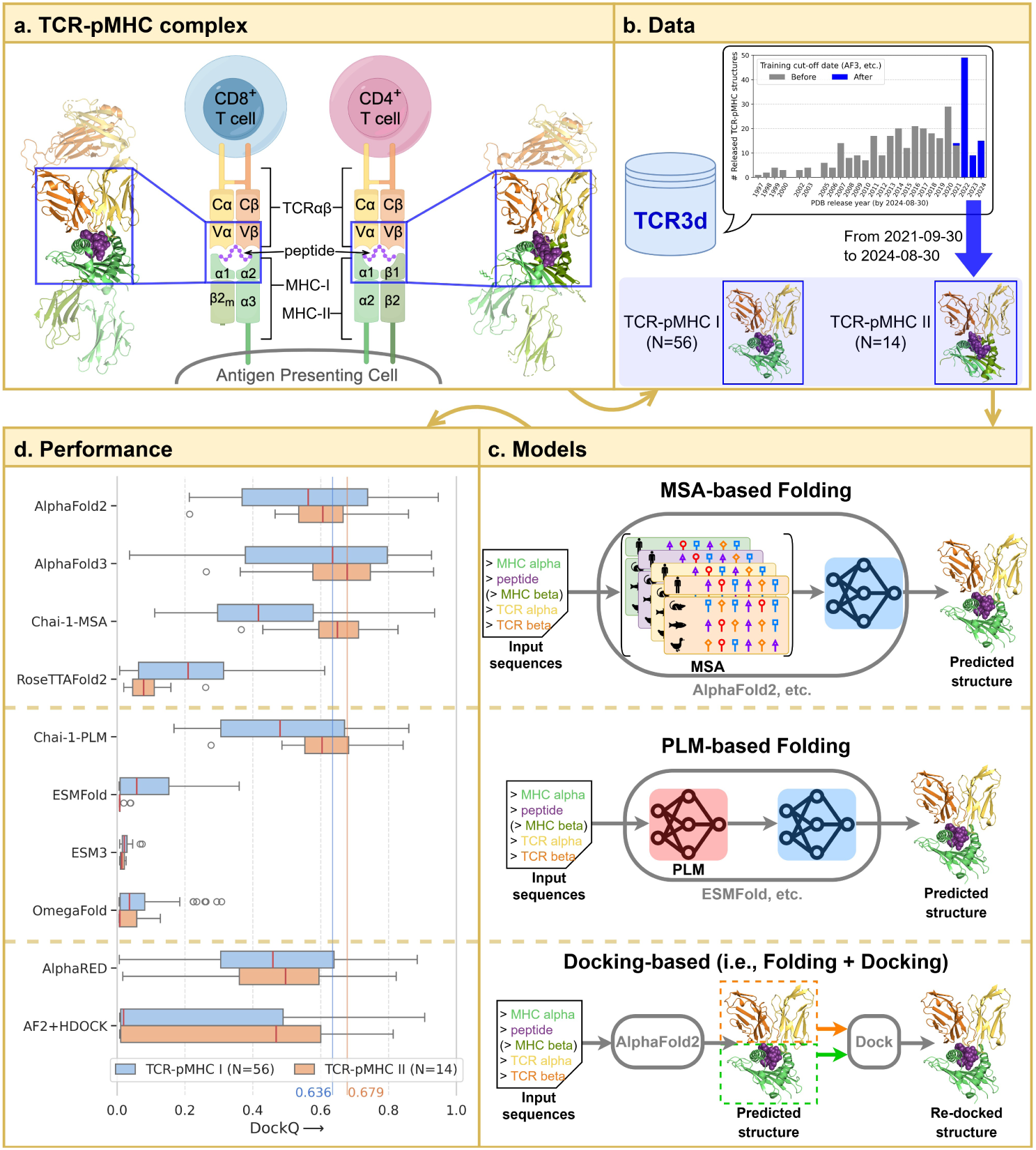
**Overview of the benchmark for TCR-pMHC complex modeling. a.** Schematic of a TCR-pMHC interaction. Key regions include the variable domains of the TCR *α* and *β* chains, the peptide, and the peptide-binding regions of class I (*α*1 and *α*2) or class II (*α*1 and *β*1) MHC. **b.** Benchmark dataset comprising 56 class I and 14 class II TCR-pMHC structures from the TCR3d database [19, 20]. All 70 structures were released after the AlphaFold3 [11] training cutoff date (2021-09-30), ensuring they were unseen during model training. **c.** Overview of three evaluated structure prediction paradigms. MSA-based methods use multiple sequence alignments to guide structure prediction via co-evolutionary signals. PLM-based methods replace MSA with protein language models for single-sequence inference. Docking-based methods refine predicted structures (e.g., from AlphaFold2 [9]) using docking algorithms to reorient the TCR and pMHC. **d.** Box-plot comparison of Top-1 DockQ [21] scores across the three structure prediction methods on the benchmark dataset. AlphaFold3 achieved the highest median DockQ scores: 0.636 for class I and 0.679 for class II TCR-pMHC complexes.

#### Superior performance of AlphaFold3

DockQ-based evaluation of overall structural docking quality is summarized in Fig. 1d. AlphaFold3 achieved the highest median DockQ scores across both Class I (0.636) and Class II (0.679) complexes, indicating that the majority of its predicted structures are of medium to high quality. Component-level modeling accuracy evaluations further revealed that AlphaFold3 generated highly accurate models for the TCR, peptide, and MHC regions, with median TM-scores exceeding 0.6 (Fig. 2d-g, Supplementary Table C1).

**Fig. 2.**
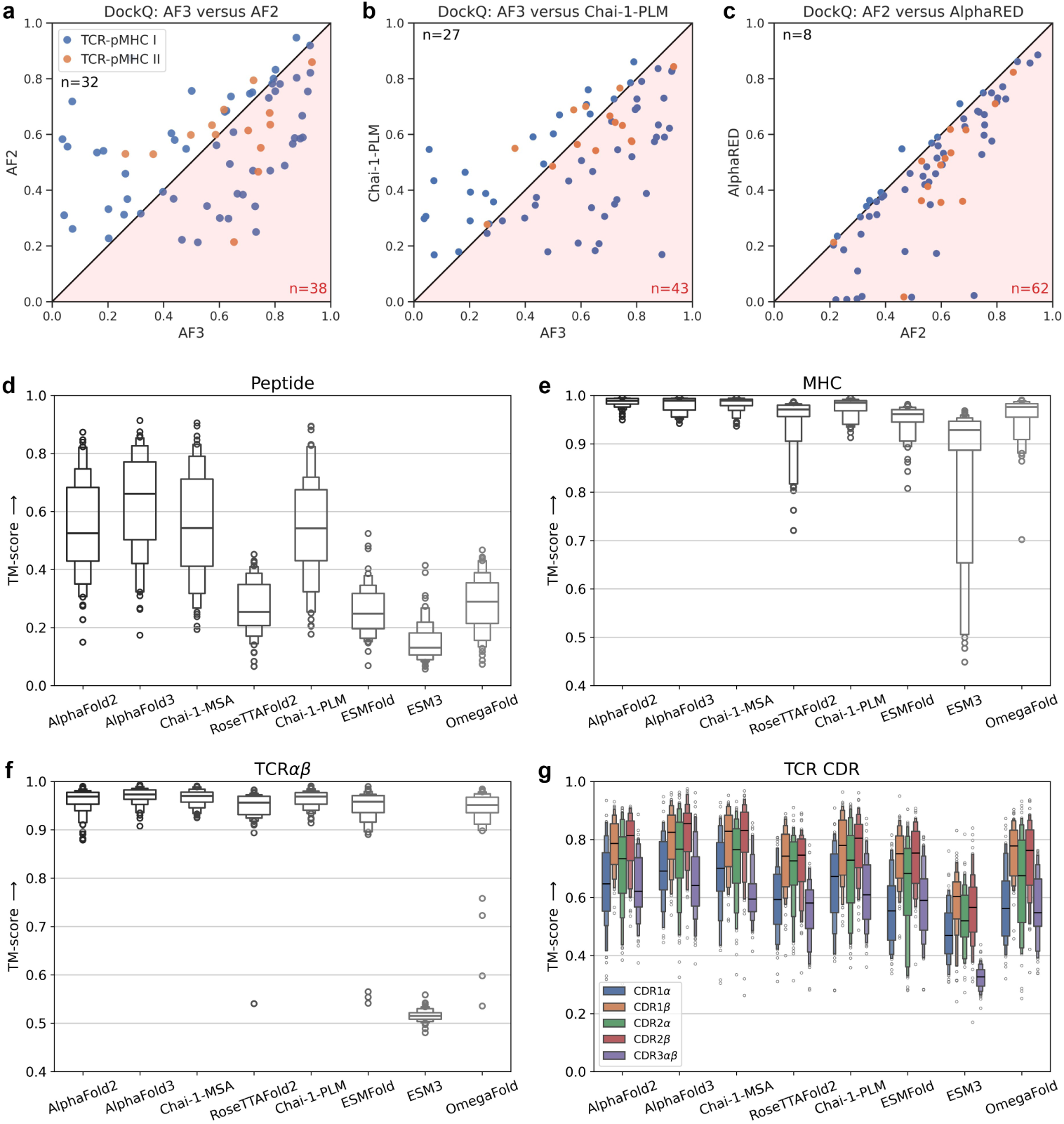
Detailed comparison of benchmark results. a.-c. DockQ scatter plots comparing Top-1 predicted structures across different models: **a.** AlphaFold3 versus AlphaFold2, **b.** AlphaFold3 versus the best PLM-based model, and **c.** AlphaFold2 versus AlphaRED. **d.-g.** TM-sore distributions of Top-1 predicted structures by MSA-based and PLM-based models for **d.** peptide, **e.** MHC, **f.** TCR, and **g.** CDR loops of the TCR.

#### Performance of MSA-based methods

Within the MSA-based models, AlphaFold2 showed slightly lower DockQ performance compared to AlphaFold3, and offered complementary predictions (Fig. 2a). In particular, AlphaFold2 occasionally generated medium-quality predictions (DockQ *>* 0.49) for complexes that AlphaFold3 failed to model correctly (DockQ *<* 0.23). Additionally, AlphaFold2 demonstrated more stable performance in MHC modeling, exhibiting lower variance than AlphaFold3. Chai-1-MSA performed comparably to AlphaFold3 on class II complexes, but lagged behind on class I complexes (Fig. 1d, Supplementary Fig. B1a). RoseTTAFold2 exhibited consistently lower performance than other MSA-based models, with reduced docking quality and notably poor peptide modeling (Fig. 2d).

While AlphaFold2 remained competitive in DockQ and structural accuracy, its routine deployment was hampered by the computationally intensive MSA generation step (taking over 120 minutes per TCR-pMHC complex in our local setup). We further evaluated two MSA-acceleration strategies based on AlphaFold2: TCR-model2 [23] with a curated TCR/MHC sequence database, and ColabFold [18] with MMseqs2 [28, 29] for rapid homology search. Both approaches yielded similar DockQ and TM-score distributions to the original AlphaFold2 (Supplementary Fig.B2a,b), while reducing MSA runtime per complex by over 70-fold (from 123.5 min to ∼1.7 min, Supplementary Fig.B2c). This highlights the practicality of AlphaFold2 when combined with optimized MSA strategies, enabling accurate and scalable TCR-pMHC structure prediction in resource-constrained settings.

#### Performance of PLM-based methods

In contrast, most PLM-based models underperformed relative to MSA-based models. ESMFold, OmegaFold, and ESM3 generally failed to produce accurate complex structures. For ESMFold and OmegaFold, the main limitation was their lack of multi-chain modeling support: both models treat multi-chain systems as single-chain inputs, resulting in incorrect inter-chain orientations. These two models produced reasonably accurate predictions for individual components such as TCR and CDR loops (Fig. 2f,g), but their peptide modeling was notably poor (Fig. 2d). In the case of ESM3, performance was further limited by its reliance on generative modeling and the use of its smallest publicly available check-point. Notably, Chai-1-PLM emerged as an exception among PLM models. It achieved performance comparable to Chai-1-MSA (Supplementary Fig. B1b) and approached AlphaFold-level quality, even outperforming AlphaFold3 in over one-third of the tested complexes (Fig. 2b). This can be attributed to Chai-1’s AlphaFold3-style architecture, which incorporates chain-aware adaptations and jointly leverages MSA and PLM features during training.

#### Performance of docking-based methods

Finally, docking-based methods failed to enhance the docking quality of the AlphaFold2-generated TCR-pMHC complexes. In AlphaRED, we performed rigid-body docking between separately predicted TCR and pMHC structures, followed by flexible refinement guided by interface pLDDT scores. However, this approach improved DockQ scores in only 8 of the 70 complexes, while performance declined in the remaining cases (Fig. 2c). A similar outcome was observed with HDOCK (Supplementary Fig. B1c), suggesting that current docking strategies offer limited added value when starting from high-quality end-to-end predictions.

### 2.2 CDR3 pLDDT: a key modeling and functional indicator for AlphaFold3

AlphaFold3 provides well-calibrated confidence scores aligned with structural accuracy and docking quality [11], serving as valuable internal indicators of model reliability. Among these, the predicted local distance difference test (pLDDT) score serves as a fine-grained metric of local confidence. Given the central role of the CDR3 loops of TCR *α* and *β* chains in antigen recognition and their inherent sequence variability [30, 31], we investigated whether the CDR3 pLDDT score could provide deeper insights into modeling quality and functional relevance. Our analysis shows that CDR3 pLDDT is not only indicative of local modeling accuracy and global docking quality (Section 2.2.1), but also useful for reranking multiple model outputs (Section 2.2.2) and detecting affinity-altering mutations (Section 2.2.3).

#### 2.2.1 CDR3 pLDDT reflects CDR3 modeling accuracy and docking quality

We first analyzed the relationship between per-residue confidence (pLDDT) and structural accuracy (RMSD) for different regions of the complex. As shown in Fig. 3a, the correlation between pLDDT and RMSD for CDR3 was found to be the most significant, with a Pearson correlation coefficient of −0.673, indicating that CDR3 pLDDT precisely reflects the accuracy of CDR3 modeling. Meanwhile, we examined the relationship of CDR3 pLDDT and docking quality (DockQ). CDR3 pLDDT correlated more strongly with DockQ than pLDDT scores from other regions of the complex or ipTM/pTM scores, which assess interface/global model quality in AlphaFold (Fig. 3b). This highlights that CDR3 pLDDT alone provides a robust local proxy for overall docking accuracy.

**Fig. 3.**
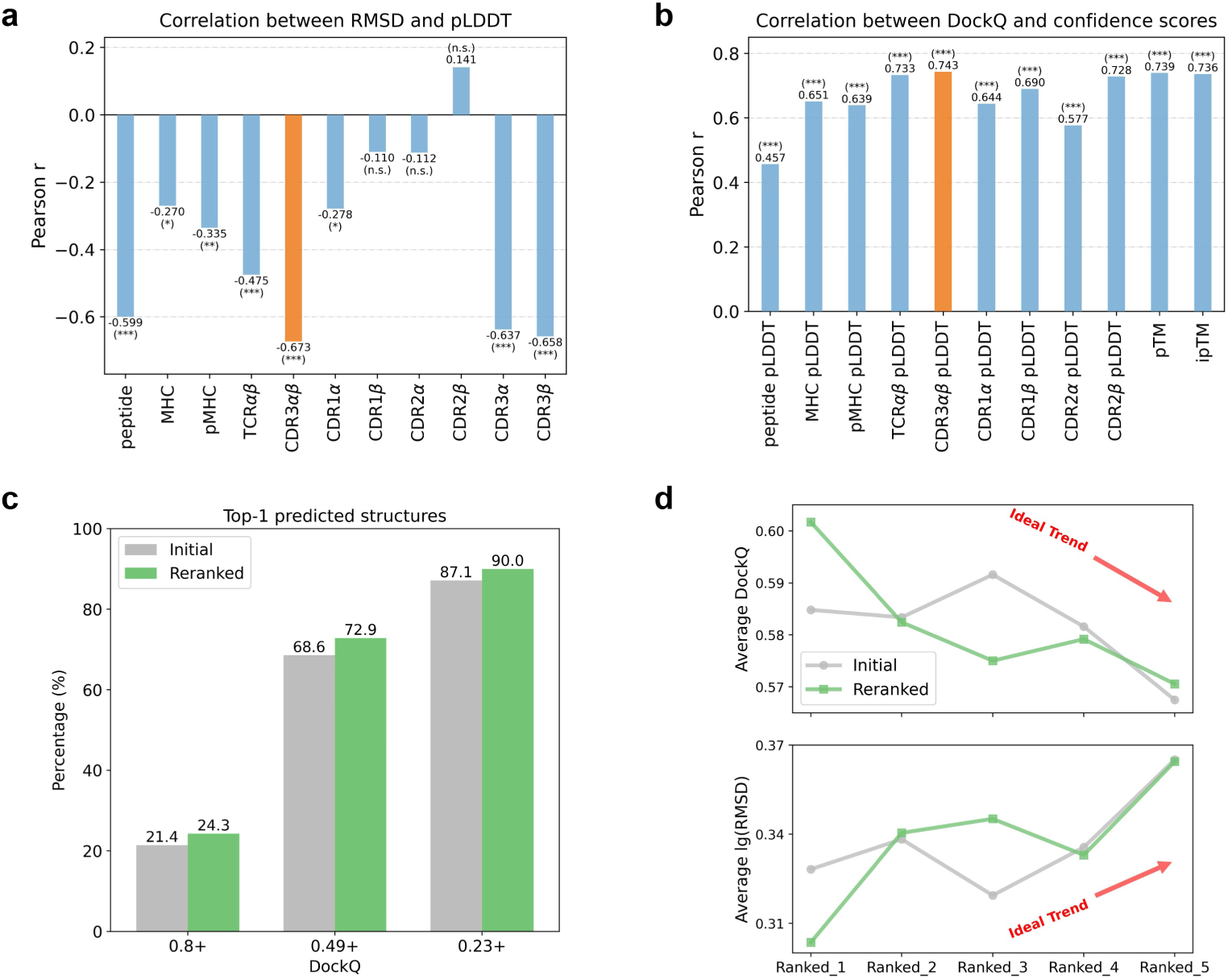
The role of CDR3 pLDDT in predicted TCR-pMHC complexes by AlphaFold3. **a.** Correlation between pLDDT scores and RMSD of different subregions. CDR3 pLDDT and CDR3 RMSD exhibit the highest correlation compared to other components. Statistical significance is indicated as: * p*<*0.05; ** p*<*0.01; *** p*<*0.001; n.s., not significant. **b.** Correlation between confidence socres (including pLDDT scores of different subregions and pTM/ipTM) and DockQ. CDR3 pLDDT also shows the strongest correlation. **c.** Comparison of DockQ distributions for Top-1 predicted structures before and after reranking AlphaFold3’s five predicted results using CDR3 pLDDT. Reranking improves the overall docking quality of Top-1 predictions. **d.** CDR3 pLDDT-based reranking enhances consistency between prediction accuracy and ranking, ensuring better alignment of docking quality with structure rankings.

#### 2.2.2 CDR3 pLDDT-based reranking enhances Top-1 structure prediction

Given the strong correlation of CDR3 pLDDT and docking quality, we reranked AlphaFold3’s five predictions using CDR3 pLDDT instead of the original ranking score. This reranking strategy effectively corrected errors introduced by the initial ranking. For the Top-1 predicted structure (Fig. 3c), reranking based on CDR3 pLDDT increased the success rate of high and medium docking quality predictions by 2.9% and 4.3%, respectively, effectively preventing the omission of high-quality predicted structures. In contrast, reranking using ipTM, pTM, AF2’s ranking score or other regional pLDDT scores yielded less improvement than CDR3 pLDDT-based reranking (Supplementary Fig. B4a). Notably, increasing the number of predictions for reranking consistently improves the reranked Top-1 quality (Supplementary Fig. B4b). Moreover, after reranking all five predictions for each complex, the docking quality and modeling accuracy showed greater consistency with the ranking (Fig. 3d), improving the reliability and confidence of the predictions. However, applying the same CDR3 pLDDT-based reranking to AlphaFold2 yielded limited improvement, with a lower correlation (∼0.4) between CDR3 pLDDT and DockQ (Supplementary Fig. B3). This may be due to AlphaFold2’s earlier confidence head, which can produce unreliable pLDDT values [32], in contrast to the refined confidence head in AlphaFold3.

#### 2.2.3 CDR3 pLDDT reflects affinity changes induced by CDR3 mutations

To further explore the functional relevance of CDR3 modeling confidence, we investigated whether CDR3 pLDDT is sensitive to binding affinity changes caused by CDR3 mutations. We used AlphaFold3 to predict the structures of both wild-type and mutant variants from the ΔΔ*G*-oriented mutation dataset, which was derived from SKEMPI v2 [33]. For each prediction, we extracted the CDR3 pLDDT score, as well as AlphaFold3’s key confidence metrics, including ranking score and interface-focused ipTM, to assess their sensitivity to affinity-altering mutations. As shown in Fig. 4a, we compared the change in confidence scores between wild-type and mutant structures for two mutations from the 1AO7 complex, one associated with increased binding affinity (ΔΔ*G <* 0) and the other with decreased affinity (ΔΔ*G >* 0). As a result, CDR3 pLDDT consistently increased or decreased in accordance with the direction of affinity change, whereas the ranking score and ipTM either stayed constant or fluctuated in ways unrelated to the affinity shift. This observation was further validated across the full mutation dataset, where CDR3 pLDDT consistently aligned with the direction of binding affinity changes more reliably than AlphaFold3’s ranking score or ipTM (Fig. 4b). Specifically, 5 out of 8 mutations associated with increased affinity (ΔΔ*G <* 0) exhibited an increase in CDR3 pLDDT; 12 out of the 13 mutations associated with decreased affinity (ΔΔ*G >* 0) showed a reduction in CDR3 pLDDT. These trends highlight CDR3 pLDDT as a more sensitive indicator of affinity-altering structural changes than AlphaFold3’s commonly used confidence scores.

**Fig. 4.**
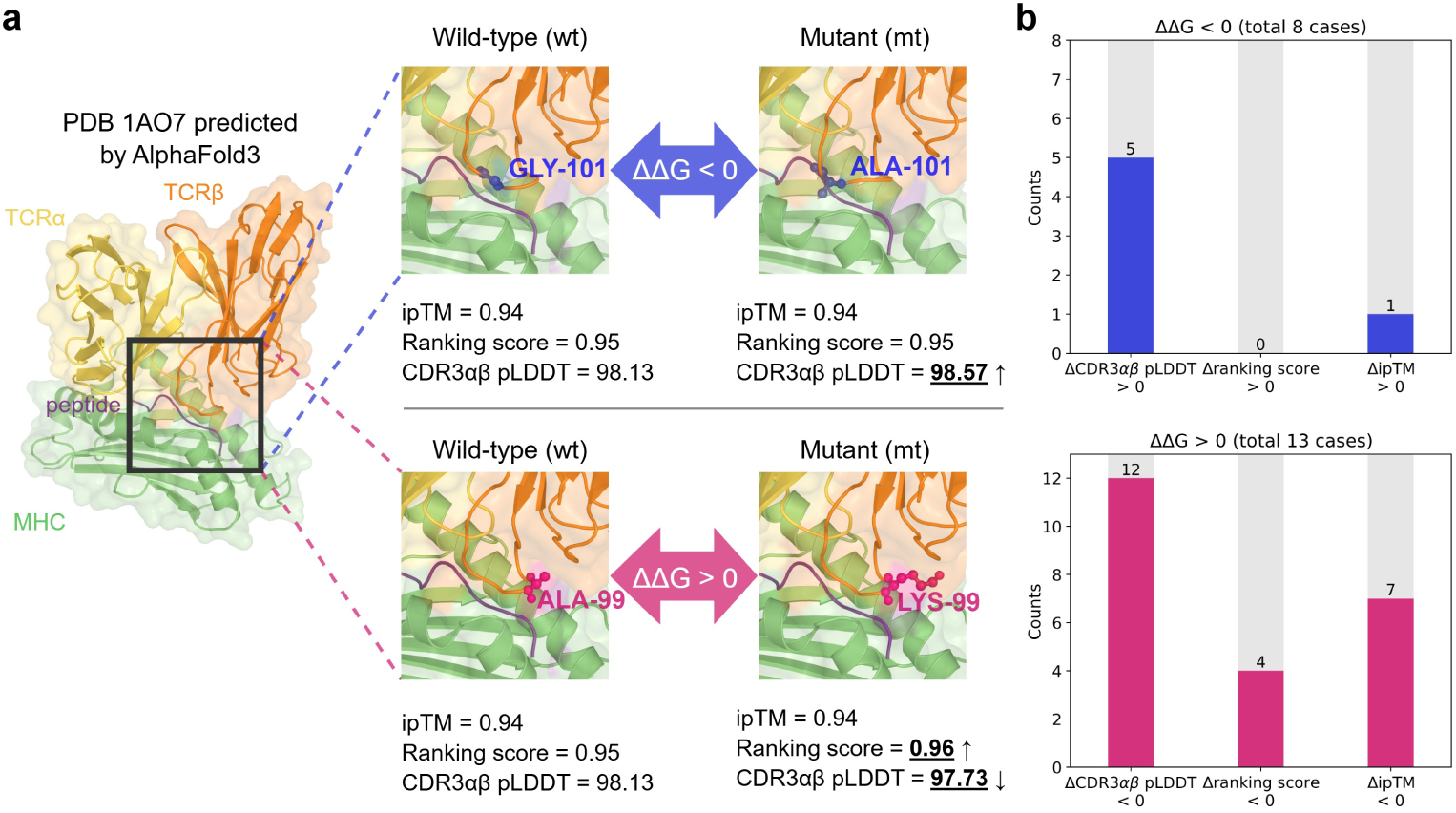
The role of CDR3 pLDDT in mutation-induced TCR-pMHC binding affinity changes. **a.** Comparison of confidence metrics for two CDR3 mutations in the 1AO7 complex with one positive and one negative ΔΔ*G*. **b.** Consistency between confidence score changes and binding affinity changes across 21 CDR3 mutations. CDR3 pLDDT changes align with the direction of affinity change in 62.5% (5/8) of ΔΔ*G <* 0 cases with increased affinity and 92.3% (12/13) of ΔΔ*G >* 0 cases with decreased affinity, outperforming ranking score and ipTM. ΔΔ*G* is defined as ΔΔ*G* = Δ*G_mutant_* − Δ*G_wild−type_*. A negative ΔΔ*G* (*<* 0) indicates that the mutation increases binding affinity, whereas a positive ΔΔ*G* (*>* 0) suggests a decrease in binding affinity.

## 3 Discussion

Using a curated set of 70 nonredundant TCR-pMHC complexes, we conducted a detailed evaluation of three prominent structure prediction methods for TCR-pMHC complex modeling, with AlphaFold [9–11] emerging as a robust performer. Our findings validated that MSA generation, a critical step in the AlphaFold pipeline, can be significantly accelerated by customizing the database [23] and employing MMseqs2 [28, 29] without loss of accuracy. This acceleration is crucial for enhancing the efficiency of AlphaFold, especially in large-scale applications where quick turnaround times are needed. Furthermore, we found that integrating prior knowledge of TCR-pMHC complexes, such as utilizing the CDR3 pLDDT as a structural confidence metric, improves the reliability of AlphaFold’s predictions and reflects mutation-induced binding affinity changes with high sensitivity. These findings underscore the value of biologically grounded insights in boosting both accuracy and interpretability in immune modeling tasks.

While general-purpose folding models like AlphaFold achieved considerable success, their adaptation to specialized systems such as TCR-pMHC complexes requires careful validation and tailored optimization. Through our comprehensive assessment, we offer actionable insights to guide the use of in-silico TCR-pMHC structure prediction in practical immunological research. These advancements are particularly important for understanding immune recognition mechanisms, which lie at the core of adaptive immunity. Moreover, accurate computational modeling of TCR-pMHC complexes will not only improve the prediction of unseen epitopes [34, 35] but also aid in the structure-based design of TCRs with higher specificity and efficacy [36, 37], especially for optimizing the CDR3 loop, which plays a pivotal role in antigen recognition. The ability to engineer TCRs with tailored specificity could significantly advance immunotherapies such as personalized cancer vaccines or TCR-based cellular therapies.

Looking forward, several promising research directions could further refine and scale in-silico TCR-pMHC modeling. First, although our results suggest that naive re-docking of predicted structures does not consistently enhance docking quality, incorporating residue-level interaction information as restraints may yield better outcomes. When integrated into docking algorithms such as HADDOCK [38], these interaction constraints could guide the modeling of binding interfaces based on biological evidence. Additionally, the potential of protein language models (PLMs) [39] for structure prediction warrants deeper exploration. Notably, our evaluation shows that Chai-1 [40] in single-sequence mode (Chai-1-PLM) achieves performance comparable to AlphaFold in modeling TCR-pMHC complexes, highlighting the promise of PLM-based folding models for delivering fast and accurate structural predictions. This opens new avenues for deploying multimer-aware PLM-based models in high-throughput neoantigen screening, accelerating the discovery of novel TCR targets. By integrating domain knowledge, advancing PLM applications, and fostering collaboration between computational and experimental research, we will move closer to realizing the full potential of in-silico structural modeling in immunotherapy.

## 4 Methods

### 4.1 Data curation

#### Benchmark dataset for structure prediction evaluation

We constructed a non-redundant benchmark dataset of TCR-pMHC complexes to evaluate model performance on previously unseen structures. The dataset was curated from the TCR3d database [19, 20], which compiles experimentally resolved TCR-pMHC complexes from the Protein Data Bank (PDB)[41]. To ensure reliable benchmarking, we applied the following filtering criteria: (1) redundancy was removed to retain unique complexes; (2) structures with a resolution higher than 3.5 Å were excluded to maintain model quality; and (3) to avoid any overlap with training data used by evaluated models, only complexes released after September 30, 2021, the latest known training cutoff across all models, were included. For consistency in model inputs, each structure was preprocessed to retain only the TCR *α*/*β* variable domains, the peptide, and the peptide-binding domains of the MHC (i.e., *α*1/*α*2 for class I and *α*1/*β*1 for class II). The final benchmark contains 56 class I and 14 class II complexes, covering 53 unique peptides, 22 distinct MHC molecules, and 51 unique TCRs, with PDB IDs listed in Appendix A.

#### ΔΔ*G*-oriented mutation dataset for confidence-function analysis

To explore the link between structural confidence and functional impact, we constructed a mutation-centered dataset for ΔΔ*G* analysis using the SKEMPI v2 database [33], which reports experimentally measured binding affinity changes in protein-protein complexes. Specifically, we selected wild-type and mutant pairs of TCR-pMHC complexes involving single-point mutations exclusively located within the TCR CDR3 region. After filtering for completeness and relevance, the final dataset comprised 21 unique mutations across 3 different TCR-pMHC complexes, each associated with an experimentally measured binding free energy change (ΔΔ*G*) (see Table C3). Among them, 8 mutations exhibited ΔΔ*G <* 0, indicating increased affinity in the mutant, while 13 showed ΔΔ*G >* 0, corresponding to reduced affinity.

### 4.2 Benchmark models

To enable a fair and comprehensive comparison across major structure prediction paradigms, we evaluated ten representative TCR-pMHC modeling tools spanning MSA-based, PLM-based, and docking-augmented approaches. For each model, we applied consistent preprocessing and postprocessing protocols where applicable, and we retained the top-ranked predicted structure per complex based on the respective ranking score for subsequent evaluation unless otherwise specified.

#### MSA-based folding models

We benchmarked four state-of-the-art MSA-based folding models: AlphaFold2 [9], AlphaFold3 [11], RoseTTAFold2 [42], and Chai-1-MSA [40]. For AlphaFold2, we used the AlphaFold-Multimer implementation from the official v2.3.2 release, setting the template database cutoff to September 30, 2021, to ensure consistency with our benchmark dataset. For each complex, we generated five candidate structures using the five available model parameter sets and selected the top-ranked prediction based on the model’s internal ranking. For AlphaFold3, we obtained predictions via the official public web server (https://alphafoldserver.com) using its default configuration. For each TCR-pMHC complex, we generated five candidate structures and selected the top-ranked prediction based on the server’s built-in scoring function. For RoseTTAFold2, we ran the model locally using the version released in January 2024. We followed the default settings provided in the official repository. For Chai-1-MSA, we generated predictions through the Chai Discovery public server (https://lab.chaidiscovery.com) in “full performance” mode, which utilizes multiple sequence alignments for enhanced accuracy.

#### PLM-based folding models

For Chai-1-PLM, we generated predictions in single-sequence mode, excluding MSAs and relying solely on the pretrained protein language model to extract sequence-level evolutionary representations. For ESM-3 [43], we used the 1.4B parameter open-source checkpoint to generate structures from the full complex sequences. For ESMFold[44] and OmegaFold[17], which do not support multi-chain inputs, we concatenated the TCR, peptide, and MHC sequences using a poly-glycine linker of length 25, following best practices from prior work [12, 45, 46]. This approach approximates inter-chain flexibility while maintaining input compatibility with single-chain models.

#### Docking-based models

For AlphaRED [47], we calculated the interface pLDDT score for the AlphaFold2 predicted structure to determine whether to perform global rigid-body docking or flexible local refinement on the receptor-ligand (i.e., TCR-pMHC) complex with ReplicaDock2 [48]. The docking structure with the highest interface energy score was then selected. Additionally, we also utilized the fast and reliable template-based docking algorithm HDOCK [49] to re-dock the predicted TCR-pMHC structures by AlphaFold2 for further evaluation. For each complex, we submitted the unbound TCR and pMHC structures as separate inputs and used the default scoring function to select the top-ranked model.

### 4.3 Evaluation metrics

#### Accuracy metrics

To assess the accuracy of predicted TCR-pMHC structures, we employed Root Mean Square Deviation (RMSD), TM-score [22], and DockQ [21] as key evaluation metrics at the C*α* atom level. RMSD quantifies the average distance between corresponding atoms in the predicted and experimentally determined structures, serving as a standard indicator of structural deviation. Prior to RMSD calculation, the predicted and native structures were aligned using the Kabsch algorithm[50] to minimize positional differences across C*α* atoms. TM-score offers a scale-independent measure of structural similarity, overcoming RMSD’s sensitivity to outliers and dependency on protein size. TM-score values range from 0 to 1, where higher scores indicate greater structural similarity, with values above 0.5 generally suggesting a correct overall fold [51]. We used both RMSD and TM-score to comprehensively assess modeling accuracy for the entire TCR-pMHC complex and its specific subregions. DockQ is a continuous metric for measuring docking model quality, combining three components: F_nat_ (the fraction of native interfacial contacts preserved in the predicted complex), LRMS (the ligand RMSD after superimposing the receptor), and iRMS (the interface RMSD between predicted and native interfacial residues). DockQ scores range from 0 to 1 and are calibrated against the Critical Assessment of Predicted Interactions (CAPRI) criteria [52, 53], classifying docking models into four categories: incorrect (DockQ *<* 0.23), acceptable (0.23 ≤ DockQ *<* 0.49), medium quality (0.49 ≤ DockQ *<* 0.80), and high quality (DockQ ≥ 0.80).

#### Confidence metrics

The predicted local-distance difference test (pLDDT) is a residue-level confidence metric proposed by AlphaFold2 [9] that quantifies prediction reliability. For specific regions of interest, such as the complementarity-determining region 3 (CDR3) loop of the TCR *α* and *β* chains, the regional pLDDT score was calculated as the average pLDDT of all residues within the region.

### 4.4 Computational environment

All models were run locally, except for AlphaFold3 and Chai-1, which were accessed through web servers. The computational setup included two Intel(R) Xeon(R) Gold 6348 CPUs at 2.60 GHz and an NVIDIA GeForce RTX 3090 GPU.

## 5 Data availability

The benchmark dataset is collected from https://tcr3d.ibbr.umd.edu/ and is avail-able at https://github.com/Jiadong001/TCR-pMHC-folding-benchmark. All structures are available in PDB (https://www.rcsb.org/). The ΔΔ*G*-oriented mutation dataset is collected from https://life.bsc.es/pid/skempi2/ and is also available at https://github.com/Jiadong001/TCR-pMHC-folding-benchmark.

## 6 Code availability

Our code to compare different methods is available at https://github.com/ Jiadong001/TCR-pMHC-folding-benchmark. The codes for different benchmark models are available via GitHub at https://github.com/google-deepmind/alphafold, https://github.com/uw-ipd/RoseTTAFold2, https://github.com/facebookresearch/ esm, https://github.com/HeliXonProtein/OmegaFold, https://github.com/ evolutionaryscale/esm, https://github.com/Graylab/AlphaRED, https://github.com/piercelab/tcrmodel2, and https://github.com/sokrypton/ColabFold.

## Acknowledgements

We thank Xiangying Pan for providing tools used in the calculation of evaluation metrics. The main cartoon components used in the schematic illustration of Fig. 1a were created with Figdraw (www.figdraw.com).

## Appendix A PDB IDs

6ZKW, 6ZKX, 6ZKY, 6ZKZ, 7DZM, 7DZN, 7L1D, 7N2N, 7N2O, 7N2P, 7N2Q, 7N2R, 7N2S, 7N4K, 7N5C, 7N5P, 7NA5, 7NDQ, 7NDT, 7NDU, 7NME, 7NMF, 7NMG, 7OW5, 7OW6, 7PB2, 7PBE, 7PHR, 7Q99, 7Q9A, 7Q9B, 7QPJ, 7R80, 7RK7, 7RM4, 7RRG, 7RTR, 7SU9, 8CX4, 8D5Q, 8DNT, 8EN8, 8ENH, 8EO8, 8F5A, 8GOM, 8GON, 8GVB, 8GVG, 8GVI, 8I5C, 8I5D, 8QFY, 8SHI, 8WTE, 8WUL, 7RDV, 7SG0, 7SG1, 7SG2, 7T2B, 7T2C, 7T2D, 7Z50, 8PJG, 8TRL, 8TRR, 8VCX, 8VCY, 8VD2

## Appendix B Supplementary Figures

**Fig. B1.**
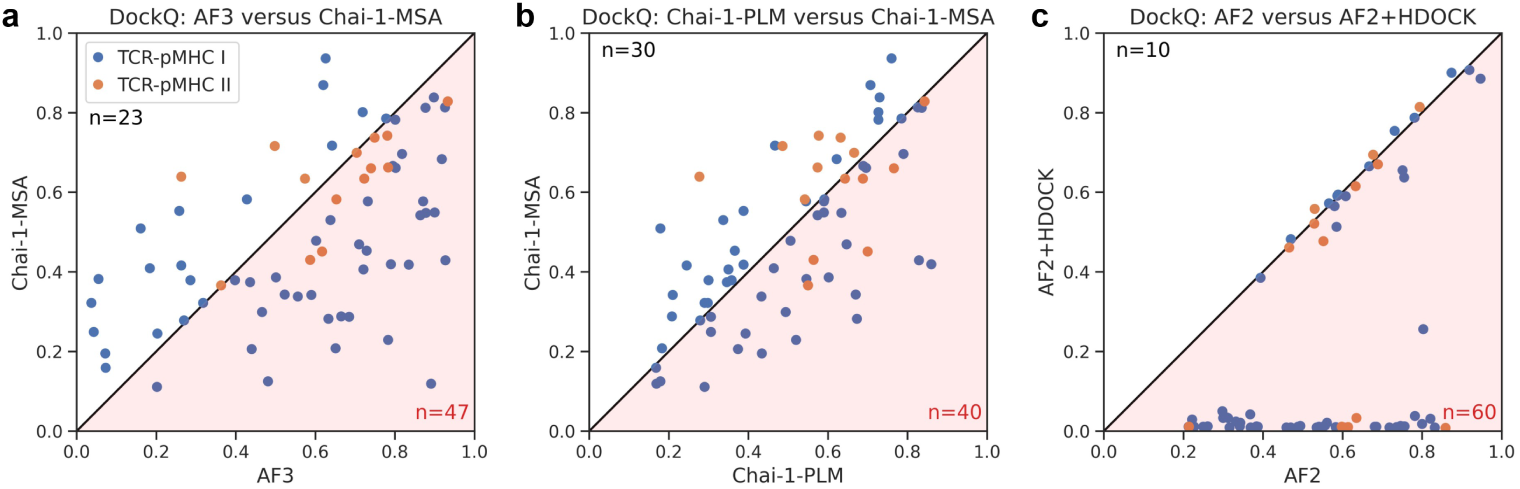
DockQ scatter plots comparing Top-1 predicted structures across different models. **a.** AlphaFold3 versus Chai-1-MSA, **b.** Chai-1-MSA versus Chai-1-PLM, and **c.** AlphaFold2 versus AlphaFold2+HDOCK.

**Fig. B2.**
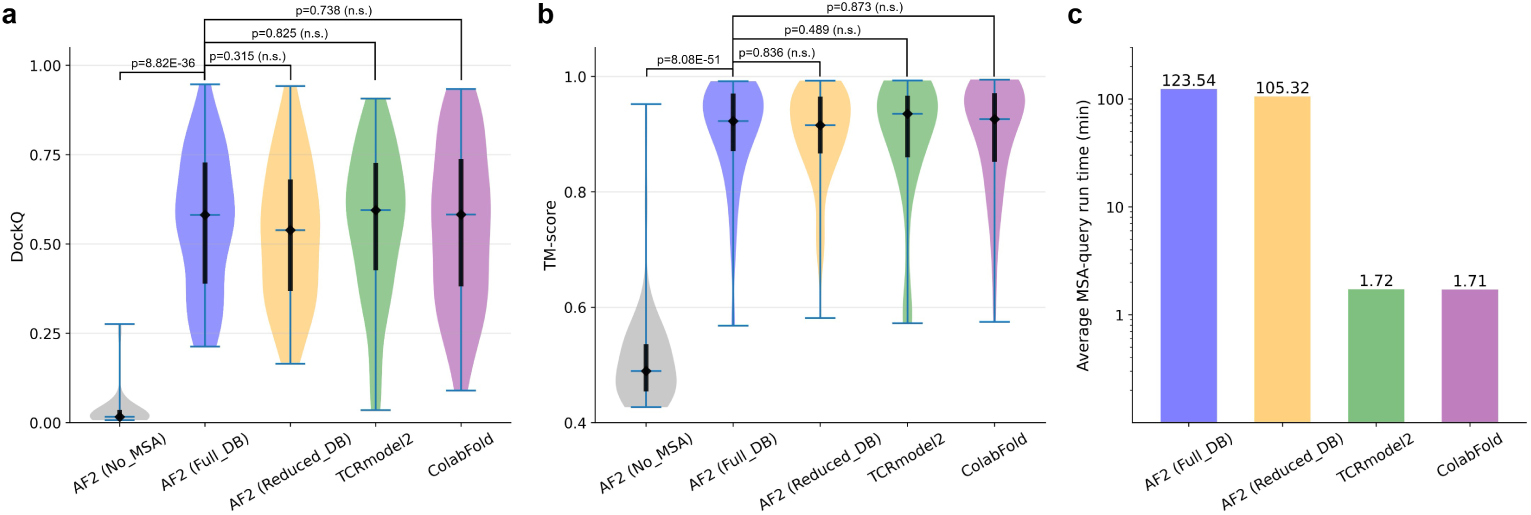
Impact of Accelerated MSA Strategies on AlphaFold2 Structure Prediction. We evaluate four MSA generation strategies for AlphaFold2: (1) AF2 (Full DB): full database search with jackhmmer and HHblits, (2) AF2 (Reduced DB): reduced database, (3) TCRmodel2 [23]: domain-specific database, and (4) ColabFold [18]: ColabFold database queried via MMseqs2 [28, 29]. **a-b.** Comparison of DockQ **(a)** and TM-score **(b)** distributions over all the 70 TCR-pMHC complexes. Naively omitting MSAs causes a significant drop in prediction accuracy. **c.** Average MSA search time per method. Both TCRmodel2 and ColabFold pipelines offer **over 70× acceleration** in MSA generation (reducing average search time to ∼1.7 minutes) compared to the full database (over 100 minutes), with negligible performance degradation.

**Fig. B3.**
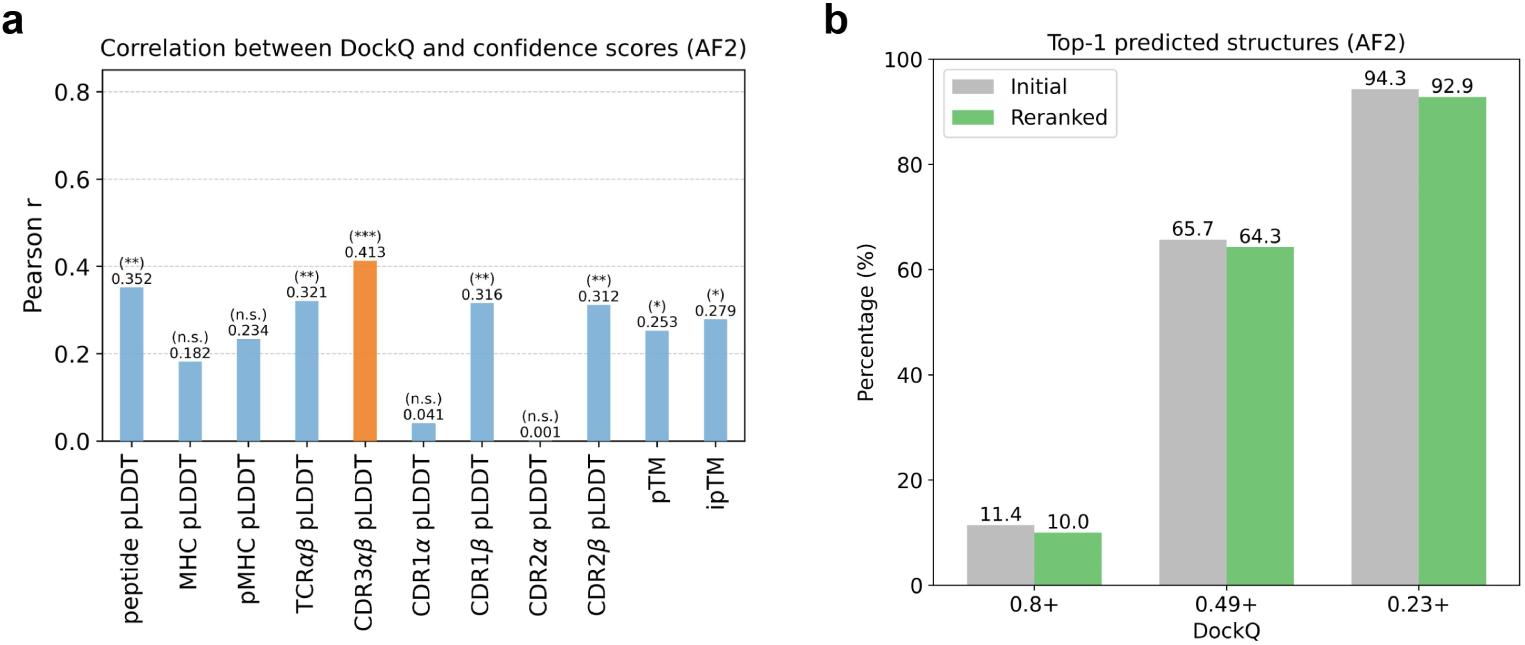
CDR3 pLDDT-based analysis for AlphaFold2. **a.** Correlation between confidence scores and DockQ. All correlation coefficients are below 0.5. CDR3 pLDDT shows the best correlation, but it is not strong. **b.** Comparison of DockQ distributions for Top-1 predicted structures before and after reranking AlphaFold2’s five predicted results using CDR3 pLDDT. Reranking with CDR3 pLDDT fails to improve the overall docking quality of the Top-1 predicted structures.

**Fig. B4.**
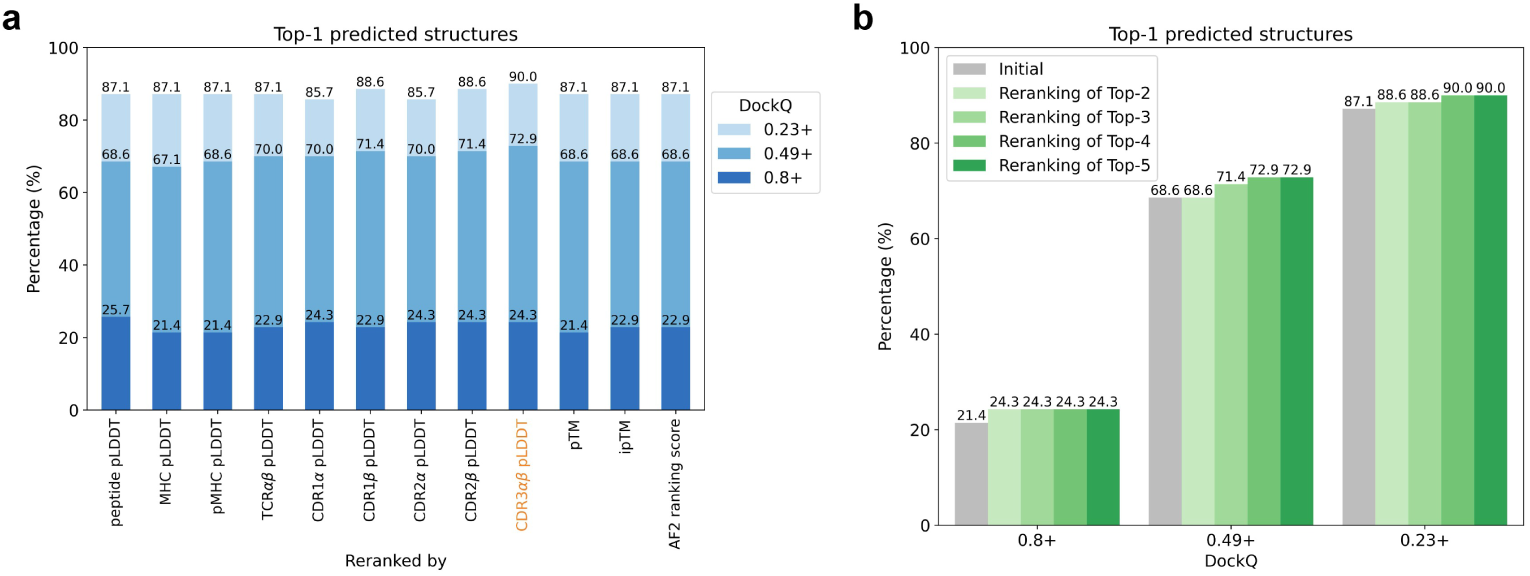
CDR3 pLDDT-based reranking improves docking quality of Top-1 predicted structures. **a.** Comparison of DockQ distributions of Top-1 predictions after reranking AlphaFold3 outputs using different confidence scores. CDR3 pLDDT-based reranking leads to the most significant improvement. **b.** DockQ distributions of Top-1 predictions after reranking the top 1–5 AlphaFold3 outputs using CDR3 pLDDT. Performance improves with larger reranking scope.

## Appendix C Supplementary Tables

**Table C1.**
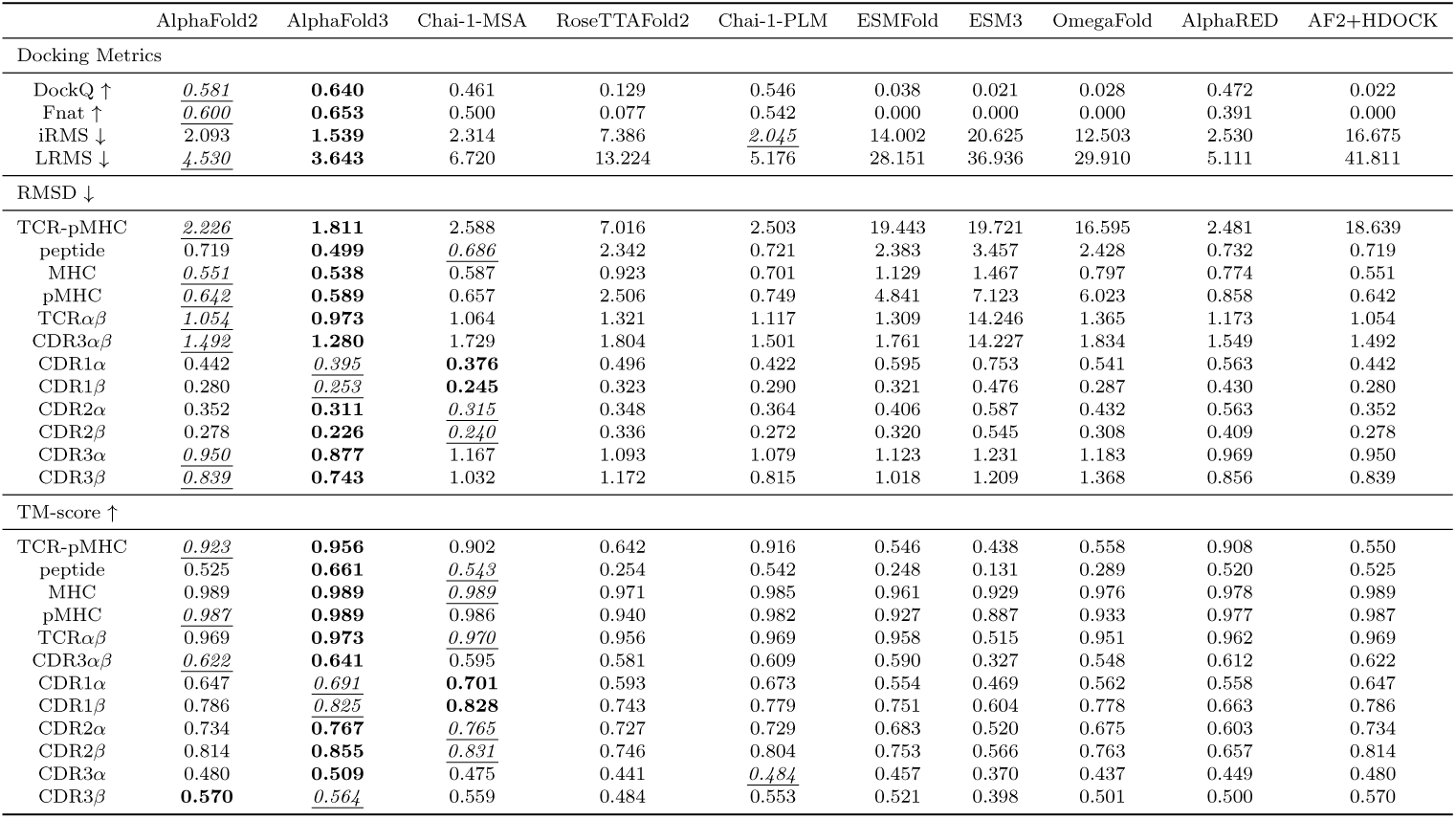
Median results of ten representative models on 70 benchmark cases. Bold and *<u>underline</u>* indicate the best and second metrics for evaluated models, respectively.

**Table C2.** Median results of AlphaFold2 with different MSA generation strategies on 70 benchmark cases. Bold and *<u>underline</u>* indicate the best and second metrics for evaluated models, respectively.

|  | AF2 (No_MSA) | AF2 (Full_DB) | AF2 (Reduced_DB) | TCRmodel2 | ColabFold |
| --- | --- | --- | --- | --- | --- |
| Docking Metrics |  |  |  |  |  |
| DockQ ↑ | 0.016 | 0.582 | 0.539 | **0.595** | <u>0.583</u> |
| Fnat ↑ | 0.000 | **0.600** | 0.552 | <u>0.575</u> | 0.571 |
| iRMS ↓ | 19.791 | 2.093 | 2.331 | **1.996** | <u>1.997</u> |
| LRMS ↓ | 44.889 | 4.531 | 5.033 | **4.453** | <u>4.472</u> |
| RMSD ↓ | | | | | |
| TCR-pMHC | 20.711 | <u>2.226</u> | 2.289 | **2.043** | 2.300 |
| peptide | 3.587 | 0.719 | 0.722 | **0.492** | <u>0.585</u> |
| MHC | 0.598 | 0.551 | <u>0.522</u> | **0.497** | 0.533 |
| pMHC | 5.916 | 0.642 | 0.671 | **0.549** | <u>0.626</u> |
| TCR$\alpha\beta$ | 5.397 | 1.054 | 1.054 | <u>0.996</u> | **0.891** |
| CDR3$\alpha\beta$ | 5.340 | 1.492 | 1.445 | **1.230** | <u>1.250</u> |
| CDR1α | 0.366 | 0.442 | 0.440 | <u>0.335</u> | **0.333** |
| CDR1β | 0.269 | 0.280 | 0.281 | **0.256** | <u>0.259</u> |
| CDR2α | 0.308 | 0.352 | 0.359 | <u>0.301</u> | **0.298** |
| CDR2β | 0.238 | 0.278 | 0.260 | <u>0.226</u> | **0.223** |
| CDR3α | 1.270 | 0.950 | 1.058 | <u>0.889</u> | **0.818** |
| CDR3β | 0.991 | 0.839 | 0.829 | <u>0.782</u> | **0.770** |
| TM-score ↑ | | | | | |
| TCR-pMHC | 0.489 | 0.923 | 0.915 | **0.935** | <u>0.926</u> |
| peptide | 0.243 | 0.525 | 0.522 | **0.625** | <u>0.606</u> |
| MHC | 0.987 | 0.989 | <u>0.990</u> | **0.991** | <u>0.990</u> |
| pMHC | 0.944 | 0.987 | 0.986 | **0.990** | <u>0.988</u> |
| TCR$\alpha\beta$ | 0.610 | 0.969 | 0.970 | <u>0.973</u> | **0.977** |
| CDR3$\alpha\beta$ | 0.410 | 0.622 | 0.614 | <u>0.655</u> | <u>0.690</u> |
| CDR1α | 0.709 | 0.647 | 0.658 | **0.752** | <u>0.735</u> |
| CDR1β | 0.801 | 0.786 | 0.794 | **0.818** | <u>0.812</u> |
| CDR2α | 0.773 | 0.734 | 0.719 | <u>0.793</u> | **0.801** |
| CDR2β | 0.839 | 0.814 | 0.825 | <u>0.849</u> | **0.853** |
| CDR3α | 0.439 | 0.480 | 0.481 | <u>0.485</u> | **0.505** |
| CDR3β | 0.542 | 0.570 | 0.557 | <u>0.573</u> | **0.582** |

**Table C3.**
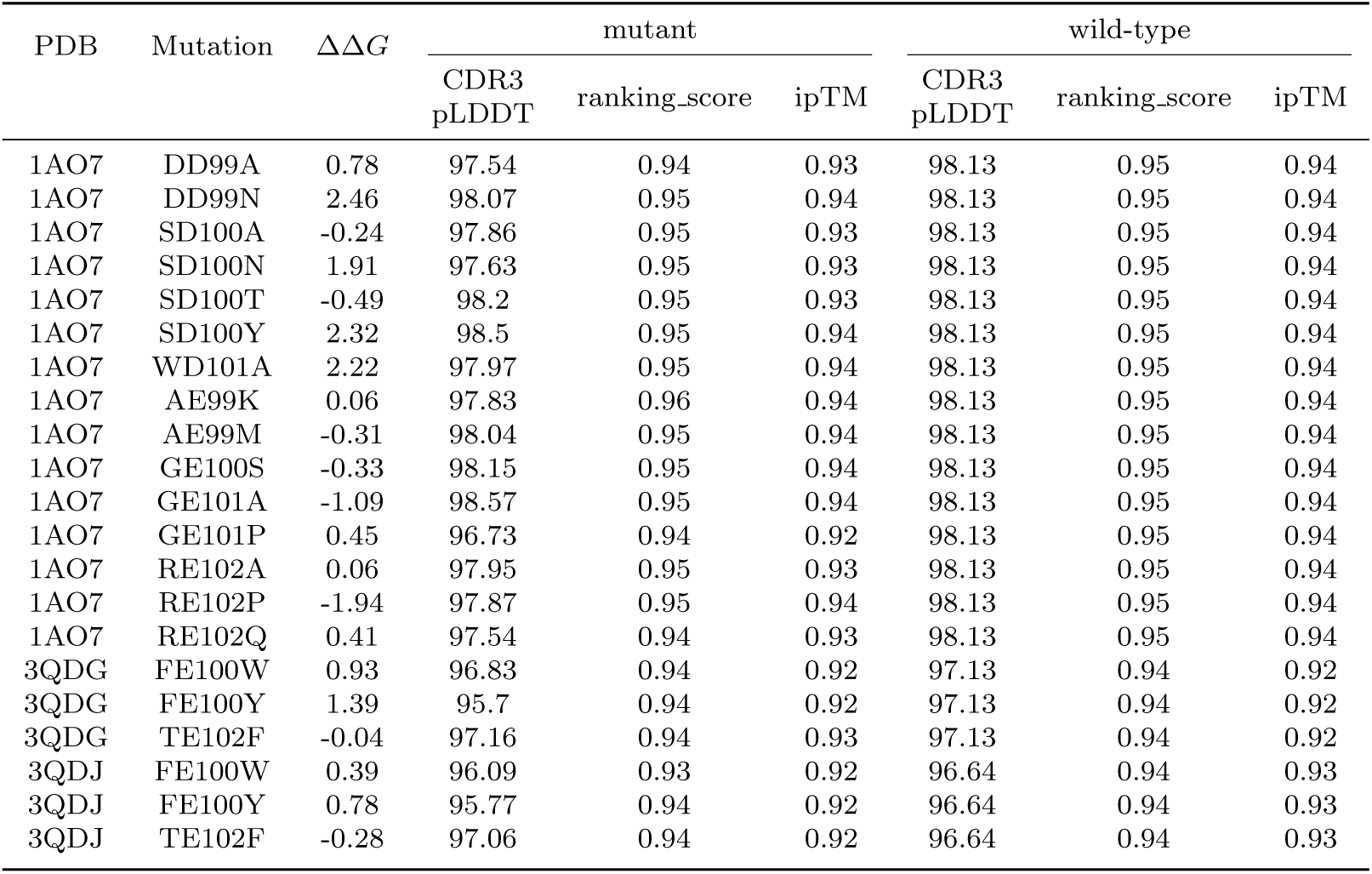
ΔΔ*G*-oriented mutation dataset details and confidence scores for wild-type and mutant structures predicted by AlphaFold3.

| PDB | Mutation | $\Delta\Delta G$ | mutant | | | wild-type | | |
| --- | --- | --- | --- | --- | --- | --- | --- | --- |
|  |  |  | CDR3<br>pLDDT | ranking_score | ipTM | CDR3<br>pLDDT | ranking_score | ipTM |
| 1AO7 | DD99A | 0.78 | 97.54 | 0.94 | 0.93 | 98.13 | 0.95 | 0.94 |
| 1AO7 | DD99N | 2.46 | 98.07 | 0.95 | 0.94 | 98.13 | 0.95 | 0.94 |
| 1AO7 | SD100A | -0.24 | 97.86 | 0.95 | 0.93 | 98.13 | 0.95 | 0.94 |
| 1AO7 | SD100N | 1.91 | 97.63 | 0.95 | 0.93 | 98.13 | 0.95 | 0.94 |
| 1AO7 | SD100T | -0.49 | 98.2 | 0.95 | 0.93 | 98.13 | 0.95 | 0.94 |
| 1AO7 | SD100Y | 2.32 | 98.5 | 0.95 | 0.94 | 98.13 | 0.95 | 0.94 |
| 1AO7 | WD101A | 2.22 | 97.97 | 0.95 | 0.94 | 98.13 | 0.95 | 0.94 |
| 1AO7 | AE99K | 0.06 | 97.83 | 0.96 | 0.94 | 98.13 | 0.95 | 0.94 |
| 1AO7 | AE99M | -0.31 | 98.04 | 0.95 | 0.94 | 98.13 | 0.95 | 0.94 |
| 1AO7 | GE100S | -0.33 | 98.15 | 0.95 | 0.94 | 98.13 | 0.95 | 0.94 |
| 1AO7 | GE101A | -1.09 | 98.57 | 0.95 | 0.94 | 98.13 | 0.95 | 0.94 |
| 1AO7 | GE101P | 0.45 | 96.73 | 0.94 | 0.92 | 98.13 | 0.95 | 0.94 |
| 1AO7 | RE102A | 0.06 | 97.95 | 0.95 | 0.93 | 98.13 | 0.95 | 0.94 |
| 1AO7 | RE102P | -1.94 | 97.87 | 0.95 | 0.94 | 98.13 | 0.95 | 0.94 |
| 1AO7 | RE102Q | 0.41 | 97.54 | 0.94 | 0.93 | 98.13 | 0.95 | 0.94 |
| 3QDG | FE100W | 0.93 | 96.83 | 0.94 | 0.92 | 97.13 | 0.94 | 0.92 |
| 3QDG | FE100Y | 1.39 | 95.7 | 0.94 | 0.92 | 97.13 | 0.94 | 0.92 |
| 3QDG | TE102F | -0.04 | 97.16 | 0.94 | 0.93 | 97.13 | 0.94 | 0.92 |
| 3QDJ | FE100W | 0.39 | 96.09 | 0.93 | 0.92 | 96.64 | 0.94 | 0.93 |
| 3QDJ | FE100Y | 0.78 | 95.77 | 0.94 | 0.92 | 96.64 | 0.94 | 0.93 |
| 3QDJ | TE102F | -0.28 | 97.06 | 0.94 | 0.92 | 96.64 | 0.94 | 0.93 |

## Notes

### Competing Interest Statement

The authors have declared no competing interest.

